# Model-based inference of gene expression noise from single-cell RNA-sequencing data

**DOI:** 10.64898/2026.06.18.733122

**Authors:** Fernanda Giersdorf, David W. Rogers, Sören Christensen, Julien Y. Dutheil

**Affiliations:** Max Planck Institute for Evolutionary Biology, Department of Theoretical Biology, Plön, Germany; Max Planck Institute for Evolutionary Biology, Department of Microbial Population Biology, Plön, Germany; Christian-Albrechts University Kiel, Department Of Mathematics, Kiel, Germany

**Keywords:** Gene expression noise, Single-cell RNA sequencing, maximum likelihood, modulated Poisson process

## Abstract

The heterogeneity of expression levels among genetically identical cells, termed gene expression noise, is a property of the gene expression process whose importance in the biology of organisms and their evolution is increasingly recognized. Measuring gene expression noise requires single-cell expression data, as obtained from single-cell RNA sequencing (scRNASeq). Its estimation, however, is challenging owing to (i) the presence of technical noise in addition to biological noise, and (ii) the heterogeneity of cell types in the sampled population. We propose a maximum-likelihood framework to infer biological noise from scRNASeq data, while accounting for technical noise, dropout probabilities, and distinct cell sequencing depths. We demonstrate the parameter identifiability using simulations and that the resulting noise estimates are uncorrelated from the mean gene expression, and therefore do not need extra correction in downstream analyses, easing intra- and inter-genome comparisons. Using two technical replicates of scRNASeq data from the wild yeast *Saccharomyces paradoxus*, we show that expression noise can be inferred in a reproducible manner.

## 1 Introduction

The molecular process of gene expression involves the interplay of many biochemical reactions leading to the synthesis of a biomolecule (typically a protein or RNA). Most of the underlying molecules, including processing enzymes, regulatory components, and DNA itself, are present in small numbers and subject to random diffusion and binding. As a result, gene expression is a stochastic process that results in a heterogeneous number of products between genetically identical cells, even when grown in a homogeneous environment [McAdams and Arkin, 1997].

Despite early recognition, the role of gene expression noise has long been understudied because it was difficult to assess experimentally. Quantitative estimation of gene expression noise has been made possible by high-throughput single-cell technologies, in particular, single-cell RNA sequencing (scRNASeq, Kim and Marioni [2013]). The variation of expression noise along the genome has been shown to correlate with several gene properties, such as essentiality [Fraser et al., 2004], protein conservation [Barroso et al., 2018], or molecular function [Newman et al., 2006], indicating that it is evolving under natural selection and therefore represents a fundamental property of gene expression, in addition to the mean expression level.

However, scRNASeq is prone to technical noise and artifacts, such as *dropouts*, where the sequencing of expressed genes fails in some cells [Kharchenko et al., 2014]. These artifacts are inherently due to the small number of molecules at play but stem from non-biological stochastic effects. Such technical noise cancels out when looking at mean gene expression, but adds up to biological noise and becomes problematic when noise is the topic of study.

The difficulty in disentangling biological from technical noise is evident in the observation that noise, even when assessed using measures based on the coefficient of variation (which is supposedly corrected for the mean), is still correlated with the mean expression level [Brennecke et al., 2013]. Single-cell RNASeq data (*D*) come as read counts per gene and per cell. The number of read counts is Poisson-distributed, proportionally to the expression level of the gene (that is, the number of messenger RNA molecules in the cell). The distribution *X* of gene expression levels across cells is therefore a latent distribution with Poisson sampling, *D* |*X ~Poisson*(*X*), which we need to infer from the observed cell counts. By the law of total variance, *V ar*(*D*) = *E*[*X*] + *V ar*(*X*), which implies *CV* ^2^(*D*) = *CV* ^2^(*X*) + 1*/E*(*X*), so that the noise and mean of the read counts are intrinsically correlated. This correlation is not biological, but stems from the Poisson process, that is, technical noise. Previous works proposed to fit a model to the noise vs. mean data [Brennecke et al., 2013] and estimated biological noise as the residual variance of this model, that is, the part of the observed variance that is not explained by the mean [Brennecke et al., 2013, Barroso et al., 2018]. However, such approaches are likely to underestimate biological variation and lead to estimates that are difficult to compare between data sets. Furthermore, raw measures of variance and mean tend to inflate noise, as they do not account for dropout probabilities.

Models of gene expression have been proposed to facilitate the analysis of single-cell gene expression. They are used primarily to perform statistical tests of differential gene expression (e.g. Kharchenko et al. [2014], Vu et al. [2016]). Here, we adapt and extend such models to estimate gene expression noise, and demonstrate their capacity to disentangle biological from theoretical noise.

## 2 Materials and methods

### 2.1 Count models in single cells

The expression profile of a gene *i* is characterized by a latent distribution *X*_*ij*_ of mRNA counts *x*_*ij*_ in different cells *j*. We aim to infer the distribution *X*_*ij*_ from the data *D*_*ij*_ = *d*_*ij*_, where *d*_*ij*_ is the number of reads for cell *j* in gene *i*.

Reverse transcriptases and polymerases will process a given sequence with fixed probability. Assuming that this probability is small and that all processing events are independent in time (no memory), the number of events that occur in a certain time interval follows a Poisson distribution with rate *λ. λ* also equals the mean and variance of the process. The number of reads in the cell *j* for the gene *i* therefore follows a Poisson distribution with a rate equal to *x*_*ij*_:

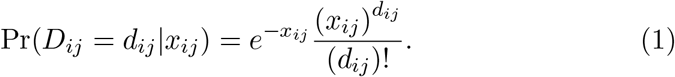

Droplet-based scRNASeq techologies use a reverse transcription step, which adds a unique molecular identifier (UMI) to the corresponding cDNA before amplification. Using UMIs counts instead of the total read counts allows to bypass amplification biases, yet not technical noise resulting from the reverse transcription. Since *X*_*ij*_ is unknown, we integrate its distribution to obtain the probability of observing *d*_*ij*_:

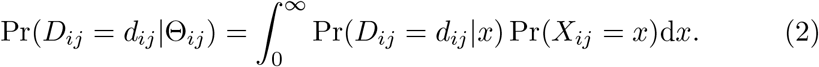

where Θ_*ij*_ is the set of parameters of the *X*_*ij*_ distribution. As *X*_*ij*_ is unknown, a common choice is to use a flexible distribution, such as a Gamma distribution (two parameters) or a scaled Beta (three parameters). As different genes have different expression profiles, they have different distribution parameters. In the simplest model, all cells share the same distribution, so that, noting *X*_*i*_ the distribution for gene *i*:

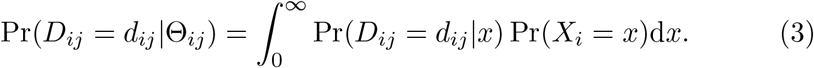

The resulting compound distributions are therefore a Gamma-modulated Poisson (equivalent to a negative binomial) and a Beta-modulated Poisson (often referred to as a BetaPoisson, Vu et al. [2016]).

Single-cell transcriptome data are characterized by additional artifacts that need to be considered. The first is heterogeneity in cell coverage: some cells yield more reads than others. This means that the polymerase has a binding probability that differs between cells. We can incorporate this into the model by introducing a cell-specific constant *l*_*j*_, so that the distribution of read counts follows a Poisson process with rate *l*_*j*_ *·x*_*ij*_. The likelihood function is then modified as

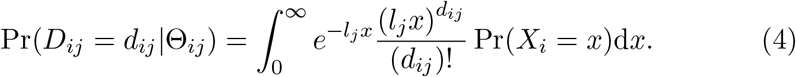

This is the approach used in Vu et al. [2016], except that their model assumes pre-normalized counts and the *l*_*j*_ parameters are introduced to account for non-integer observations. Here, we directly model the raw data and use *l*_*j*_ as a normalization constant.

Next, we need to account for the dropout probability. A dropout event occurs when transcripts fail to be amplified [Kharchenko et al., 2014]. This leads to zero-inflated distributions of read counts, which is modeled as a mixture distribution:

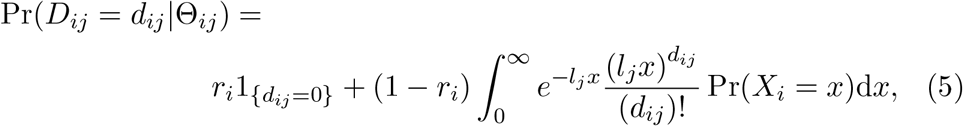

where *r*_*i*_ is the dropout probability, which we model as a gene-specific property. This model is equivalent to the five-parameter BetaPoisson model of Vu et al. [2016].

The integral in the likelihood calculation cannot be computed analytically and requires numerical integration. Instead, we discretize *X*_*i*_ into *k* = 1..*t* categories, each with an average value *x*_*ik*_ and a probability *p*_*ik*_. This leads to the simplified likelihood formula:

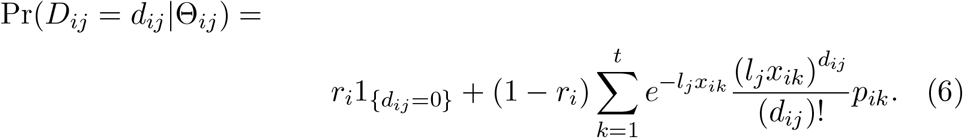

The likelihood of the data is then simply given by the product of all gene-cell likelihoods:

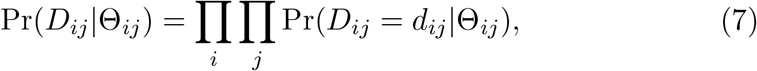

where Θ_*ij*_ includes the sets of *l*_*j*_ and *r*_*i*_, as well as the gene specific parameters of the distribution of gene expression.

### 2.2 Measures of gene expression noise

In this framework, technical noise is modeled by the Poisson process. Biological noise is derived from the distribution of gene expression *X*. We first consider a Gamma distribution with shape parameter *α* and rate parameter *β*. For convenience, we consider a reparametrization where *β* = *α*, and introduce a scaling parameter *γ*, so that the probability density function becomes

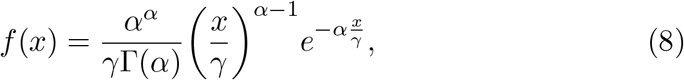

where Γ(*x*) is the *Gamma* function 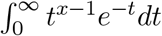. The Gamma distribution has mean *µ* = *γ* and variance 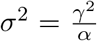. The noise of the distribution, *η*, is defined as the square of the coefficient of variation, a dimensionless value equal to the variance divided by the square of the mean. Each gene *i* has its own distribution with parameters *γ*_*i*_ and *α*_*i*_, so that:

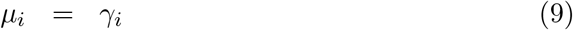

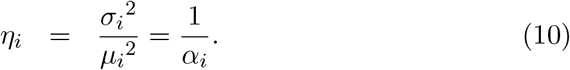

In the case of a scaled beta distribution with shape parameters *α* and *β*, and a scaling factor *γ*, the probability density function is

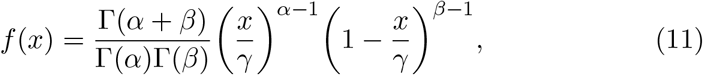

with mean 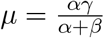 and variance 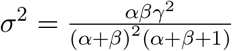. For each gene *i* with a distinct Beta distribution with parameters *α*_*i*_, *β*_*i*_, and *γ*_*i*_. The mean and noise of the scale beta distribution can then be written as

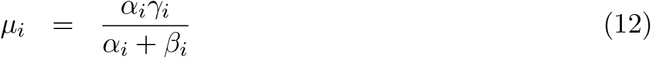

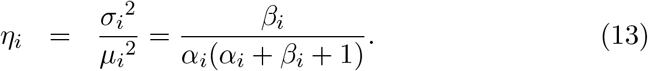

### 2.3 Cell partition models

To account for cell heterogeneity within a sample, we can extend the model to allow different sets of cells to have distinct distributions. We define a partition *C* with *q* clusters of cells *C*_*q*_ = {*c*_*j*_ = *x*}, where *x ∈ *1..*q*. Equation 6 becomes:

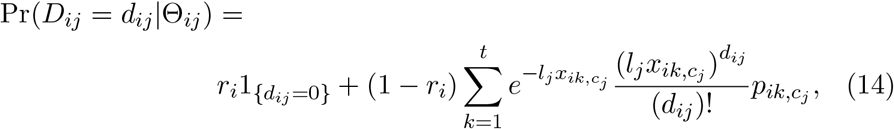

where {*x*_*ik,c*_, *p*_*ik,c*_} denotes the value and probability of the *k*th category of the distribution for gene *i* in cell *j*. Each cell is assigned a distribution from a given set. As cell type clusters are typically determined solely based on the mean expression levels, it is also possible to force distributions to share parameters. A sensible choice consists of allowing different scale parameters between cell groups, but shared shape parameters. However, in principle, the model can be used to test whether different groups of cells have different levels of noise.

Partition models can be compared using Akaike’s Information Criterion (AIC). In case partition models are nested, a likelihood ratio test (LRT) is also possible, allowing the assessment of the significance of a certain set of cell clusters, compared to a more coarse clustering. Similar tests are possible (although approximate) on a per gene basis, using profile likelihoods instead of the full likelihood (see below).

### 2.4 Parameter estimation

Parameters can be estimated by maximizing the log likelihood function, to avoid numerical underflow resulting from the multiplication of many probabilities. However, the total number of parameters is large: for a *n ×m* data matrix of *n* genes and *m* cells, we have *m* normalization constant *l*_*j* ∈ 1..*m*_, *n* dropout probabilities *r*_*i∈* 1..*n*_, and *n× z* gene specific distribution parameters, where *z* = 2 for the Gamma distribution and *z* = 3 for the scaled Beta distribution (*z* being further multiplied by the number of clusters, in case partition model is used). For computational efficiency, we maximize the profile likelihoods Pr(*D*_*i·*_|Θ_*i·*_) for each gene *i*, where *D*_*i·*_ = *{d*_*ij*_*}*_*j∈*1..*m*_ and 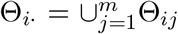, allowing the estimation of gene-specific parameters independently for each gene:

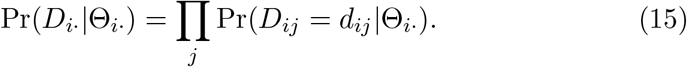

We can do the same for the cell-specific normalization constant by maximizing the corresponding profile likelihood of each cell *j*, Pr(*D*_*·j*_|Θ_*·j*_):

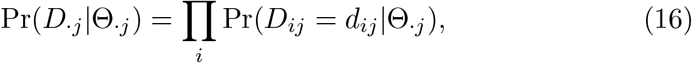

where *D*_*·j*_ = *{d*_*ij*_*}* _*i∈*1..*n*_ and 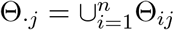.

Furthermore, we demonstrate that under asymptotic conditions (i.e., many genes with no single gene dominating the counts in each cell) the library-size estimator

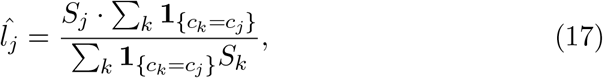

is a consistent estimator of the normalization constants *l*_*j*_, where *S*_*j*_ = ∑_*i*_ *d*_*ij*_ is the total number of counts of all genes *i* in cell *j* (see Asymptotic estimates of cell normalization parameters). By analogy with pooled RNA sequencing, *S*_*j*_ would be the “library size” for the sample *j*. The estimator 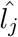 is then equal to the cell library size divided by the average library size of the cells in the group to which the cell *j* belongs. When no partition is specified (that is, all cells belong to the same cluster), the denominator is then the average cell library size in the full sample.

### 2.5 Model implementation

The model was implemented in a standalone program written in C++20, called `N.I.S.C.E.` (Noise Inference from Single-Cell Expression data). The C++20 specification level allows the use of built-in multi-threading to speed up likelihood calculations and parameter optimization. At the time of writing, the code can be compiled using the gcc compiler (version *≥*12), or any compiler with support for `execution::par. N.I.S.C.E.` also requires the libraries `boost::math_c99, TBB, Zlib and Bio++ bpp-core` (version *≥* 3.00) [Guéguen et al., 2013].

### 2.6 Simulations

Simulated datasets were obtained by drawing random numbers from zero-inflated Gamma (resp. scaled Beta) modulated Poisson distributions for 5,000 genes and 1,000 cells. Cell normalization constants were randomly chosen for each cell from a uniform distribution between 0.5 and 1.5. Gene-specific dropout probabilities were randomly chosen from a uniform distribution between 0 and 1. The gene distribution parameters *α, β*, and *γ* were taken randomly from uniform distributions between 0 and 100.

### 2.7 Single-cell data generation

The yeast strain *Saccharomyces paradoxus* CBS8841 was purchased as a lyophilized culture from the Westerdijk Fungal Biodiversity Institute’s fungal and yeast collection (Centraalbureau voor Schimmelcultures – CBS) and grown in nutrient-rich, high-glucose GPY medium at 20°C, harvested in exponential growth phase (OD600 rep1: 0.34, rep2: 0.32, measured using Ultraspec 10 Cell Density Meter (Biochrom, Art.-Nr. 10704417).

The samples were processed using a 10x Genomics Chromium protocol adapted to digest the yeast cell wall by enzymatic protoplasting, ensuring efficient cell lysis [Vermeersch et al., 2022]. In summary, a zymolyase solution was prepared by dissolving 2,000 U of Zymolyase Ultra (Zymo Research, Art.-Nr. E1007-2) in 285.7 µL of 1*×* Quantiscript RT buffer (Qiagen QuantiTect Rev. Transcription Kit, Art. Nr. 205311). The solution was sterilized using a 0.22 µm centrifugal filter spun at 12,000 *×* g for 1 minute (Sigma Aldrich, Art.-Nr. UFC30GV0S). GEM and libraries were constructed using the Chromium Next GEM Single Cell 3’ Kit v3.1 (10x Genomics, PN-1000268), following the user guide (CG000315 [10x genomics, 2024]) with the modification for the lysis of fungal cells from [Vermeersch et al., 2022]: The Gel Beads Kit v3.1 (PN-1000129) was thawed, 1 µL of gel beads was removed from each tube and replaced with 1 µL of the zymolyase solution. The tubes were resealed with sequencing-plate tape, vortexed for 30 seconds, and briefly centrifuged (5 seconds). The beads were then loaded onto the Chromium Chip G following the standard v3.1 protocol.

The library concentration was quantified using the Qubit dsDNA HS Assay Kit (Invitrogen, Thermo Fisher Scientific) on a Qubit 3.0 Fluorometer (Catalog number Q33216). Sequencing was performed on the Illumina NovaSeq 6000 platform (paired-end, 100 bp), targeting a sequencing depth of 30,000 reads per cell, following 10x Genomics guidelines.

The genomic sequence and annotation were acquired from the Yeast PacBio Reference Panel (YPRP), where the CBS8841 strain is named UFRJ50816 [Yue et al., 2017]. The GFF3 annotations were converted to the GTF format using gffread [Pertea and Pertea, 2020], excluding tRNAs. Custom genomic references were generated using the function mkref in Cell Ranger (9.0.0) [Zheng et al., 2017]. Single-cell gene expression matrices were made using the function count from Cell Ranger, which returns numbers of unique molecular identifiers (UMI) for every gene in every cell. Downstream processing was performed using Seurat [Hao et al., 2023]. For each sample, a minimum threshold of 200 total RNA counts per cell was applied to remove low-quality barcodes. To remove potential doublets, an upper bound was set for each sample, defined as the median total RNA count plus three times the median absolute deviation of the sample. The data were normalized, scaled, and cell clustering was performed using the function FindClusters.

To optimize the partitioning of cell states, cells were divided into 1 to 8 clusters per sample by varying the resolution parameter.

### 2.8 Statistics

#### 2.8.1 Correlation of measures between technical replicates

To remove the effect of the number of cells with no observations (zero-counts), we implemented a partial correlation approach. Given that a linear model between the variable of interest and the number of observations did not provide a good fit, we employed generalized additive models (GAM). The models were fitted using the mgcv package (version 1.9-4) in R [Wood, 2011].

#### 2.8.2 Gene set enrichment analyses

Gene set enrichment analyzes were conducted using the `fgsea` package [Korotkevich et al., 2019], using the Gene Ontology annotation from the `org.Sc.sgd.db` package [Carlson, 2025] and KEGG Pathway annotations for the `KEGGREST` package [Tenenbaum, 2025]. Genes were ranked according to their noise levels, as estimated in several models with increasing numbers of clusters. GO terms and KEGG pathways significant at the 1% in at least one model were reported.

## 3 Results and discussion

### 3.1 Expression distribution parameters are identifiable

We simulated data under the models introduced above in order to assess whether the parameters are identifiable. A data set of 5,000 genes in 1,000 cells was drawn from independent gene distributions with random parameters (see Materials and methods). Parameters were then re-estimated by maximum likelihood using the model used for simulating the data. We also compared with estimates from the simpler Gamma model when the data were generated under the more parameter-rich scaled Beta model.

Cell-specific normalization parameters were estimated by maximum likelihood and are perfectly recovered under all conditions (Figure 1A). We further show that the asymptotic forms give a correct estimate of the norm (Supplementary Figure S1). The dropout probabilities are also well recovered, although with a larger estimation variance (Figure 1B). When data were simulated under a scaled Beta model, the estimation of dropout parameters hit the zero limit more often, even when the true value was high (Figure 1B, second and third columns).

**Figure 1:**
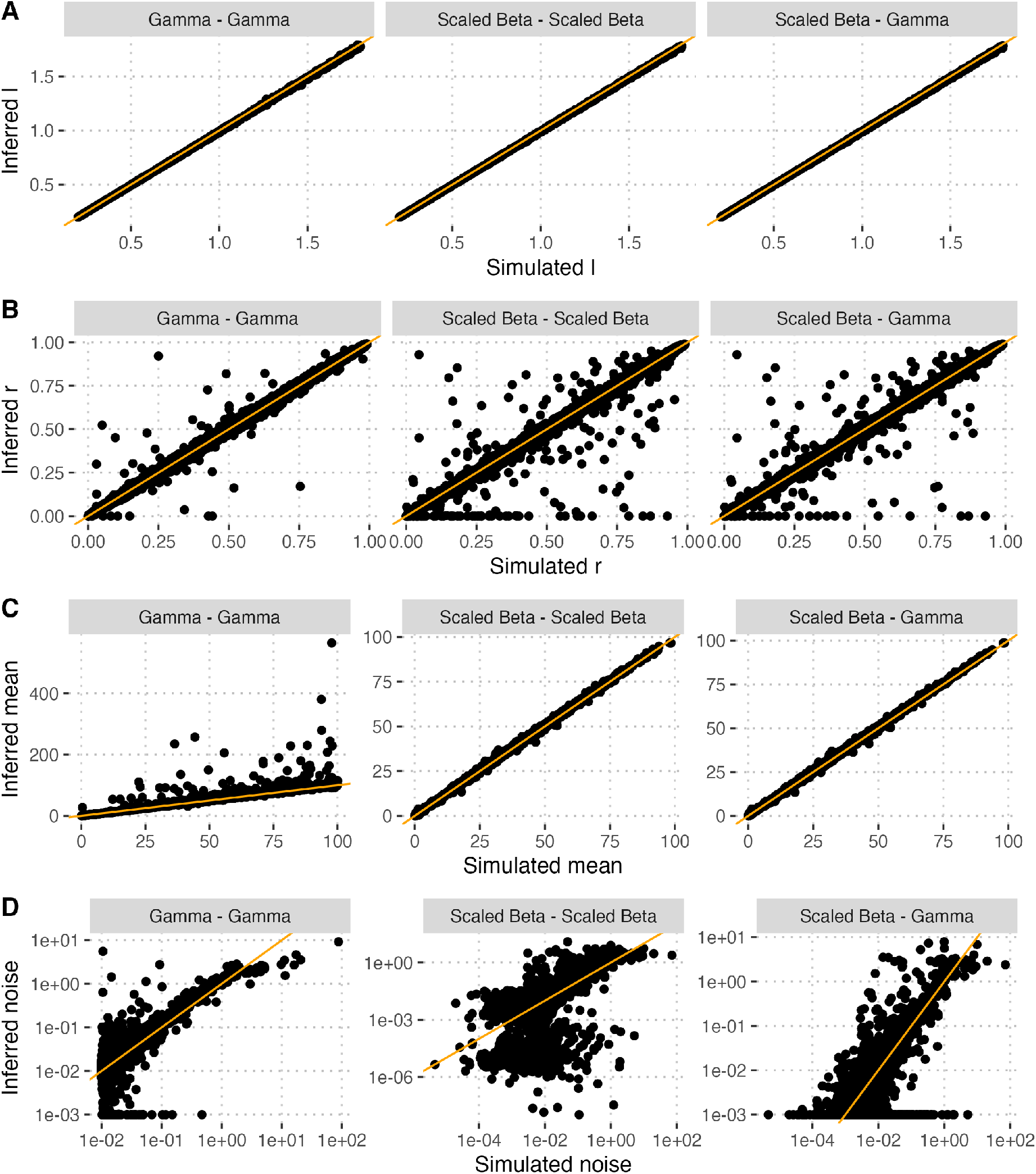
Parameter identifiability. Simulated datasets of 5,000 genes in 1,000 cells from random sets of parameters, using a Gamma (first column) or a scaled Beta (second and third columns) model. Parameters and re-estimated using a Gamma (first and third columns) or scaled Beta (second column) model. A) Normalization constants, *l*. B) Dropout probability, *r*. C) Mean gene expression, *µ*. D) Gene expression noise, *η*.

Mean gene expression values were generally well recovered, but with a lower variance when data were simulated under a scaled Beta model (Figure 1C). Gene expression noise was the most difficult quantity to recover. The estimation is unbiased when a Gamma model is used for inference, even though the data were simulated under a scaled Beta model. The two models differ by two important aspects that may explain these differences in behavior: 1) the scaled Beta distribution is bounded between 0 and the scale parameter *γ*, while the Gamma distribution is defined on [0, +*∞* [; 2) in the Gamma distribution, the mean and noise are independent, while they are negatively correlated in the scaled Beta distribution.

Based on these simulation results, we conclude that the scaled Beta model is to be favored when the focus of the inference is the mean expression, as it provides measures with lower estimation variance. However, the Gamma model performs globally better, at least for data sets in the size range of the one tested here: it provided unbiased estimates of the mean expression and expression noise, even when the underlying data are not Gamma-distributed.

### 3.2 Model-based inference can distinguish biological from technical noise

As a mean of comparison, we also computed classical estimates of mean and noise on the simulated data. The data is normalized for the total read counts per cell, following standard practices:

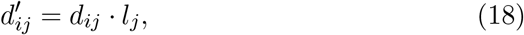

where *l*_*j*_ are computed using equation 17. The expression mean and noise are then directly computed from the normalized data:

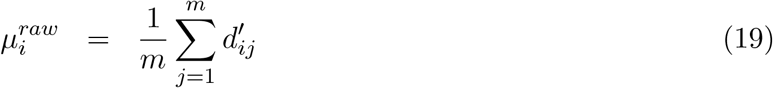

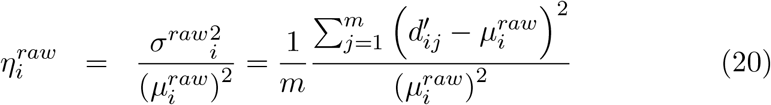

and similar estimates after discarding all counts equal to zero (hereby referred to as “star” estimates):

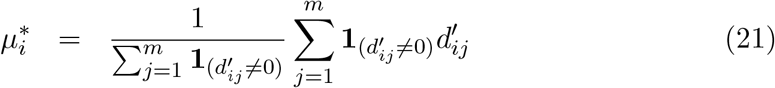

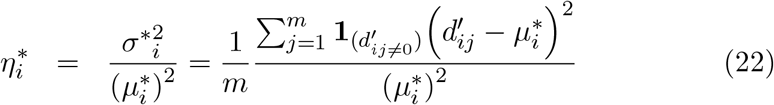

Raw counts only provide an upper bound to the true mean expression rate, while non-zero estimates provide an unbiased estimation (Figure S3A). Conversely, raw counts only provide a lower bound of the expression noise, and non-zero estimates are still largely overestimated (S3B). Furthermore, as expected, raw and non-zero counts lead to strong negative correlations between the estimated mean and noise, even when the true values are uncorrelated (Table 1).

**Table 1:** Correlation between expression mean and noise.

| **Variables** | **Correlation<sup>a</sup>** | **P values** |
| --- | --- | --- |
| Simulated<sup>b</sup> | -0.0050 | 0.5925 |
| Inferred, raw estimates | -0.5474 | < 2.2e-16 |
| Inferred, non-zero estimates | -0.5166 | < 2.2e-16 |
| Inferred, model-based estimates<sup>1</sup> | 0.0032 | 0.7354 |

<sup>a</sup> Kendall's correlation of ranks
<sup>b</sup> Data were simulated under a Gamma model with independent mean and shape.

### 3.3 Biological noise estimates are repeatable

We inferred expression noise from single-seq RNA sequencing data obtained from cultures of the yeast *Saccharomyces paradoxus*, a wild relative of the baker’s yeast *S. cerevisiae*. We conducted two replicate experiments under the same conditions and estimated the mean expression and expression noise after estimating the parameters using a Gamma model (see equation 9) and compared the estimates for all genes between the two replicates (Figure 2). The estimates of mean expression appear to be highly repeatable between the two replicates (Figure 2A). The noise estimates are highly correlated between replicates (Kendall’s *τ* = 0.49, P value < 2.2e-16), but show several uncorrelated estimation failures, where noise is estimated to the minimal bound in one of the two replicates. Estimation failures generally involve genes with low cell counts, but not exclusively (Figure 2C).

**Figure 2:**
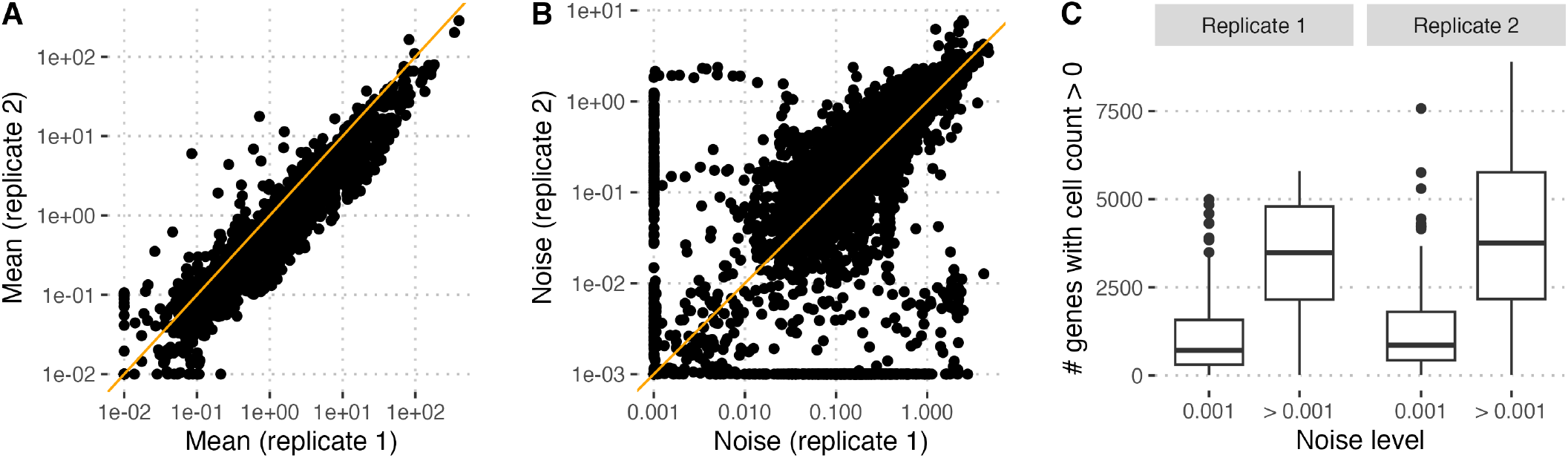
Estimation of gene expression mean and noise under a Gamma model for two technical replicates. A and B: estimates of replicate 2 plotted against estimates of replicate 1, for expression mean (A) and noise (B). Each point represents one gene. C: number of cells with observations plotted as a function of noise estimation failure (Noise estimate equal to the lower bound, 0.001).

To assess whether model-based inference effectively disentangles biological from technical noise, we looked at the strength of the correlation of different statistics between technical replicates (Figure 3). We expect that model-based estimators, which measure biological noise only, should be more strongly correlated than raw statistics, which measure a combination of biological and technical noise. To further assess the impact of the amount of available data, we look at the correlation between replicates after discarding genes with observations (non-zero counts) in less than a given number of cells. We show that model-based estimates only lead to a modest improvement in the estimation of mean expression (Figure 3A), which is expected since technical noise should only affect the variance of the mean estimation. However, the effect was very strong for noise estimation (Figure 3B). Since the number of observations was highly correlated between the two replicates (Figure 3C), we used partial correlations to account for this spurious correlation (see Materials and methods). Figure 3B shows that model-based noise estimates are more correlated between replicates than simple statistics, demonstrating their ability to account for technical noise. All methods, but Gamma-based estimates show increasing correlations as genes with less observations are removed. Although the Gamma model leads to the highest correlation for genes with fewer observations, its effectiveness decreases as the number of observations increases. We posit that this is due to the Gamma distribution not being a perfect fit to the real data, the scaled Beta model being a better fit, owing to the presence of an additional parameter. However, the addition of this extra parameter comes at the cost of identifiability: consistent with the simulations, the model can be fitted in fewer genes (Figure 3D).

**Figure 3:**
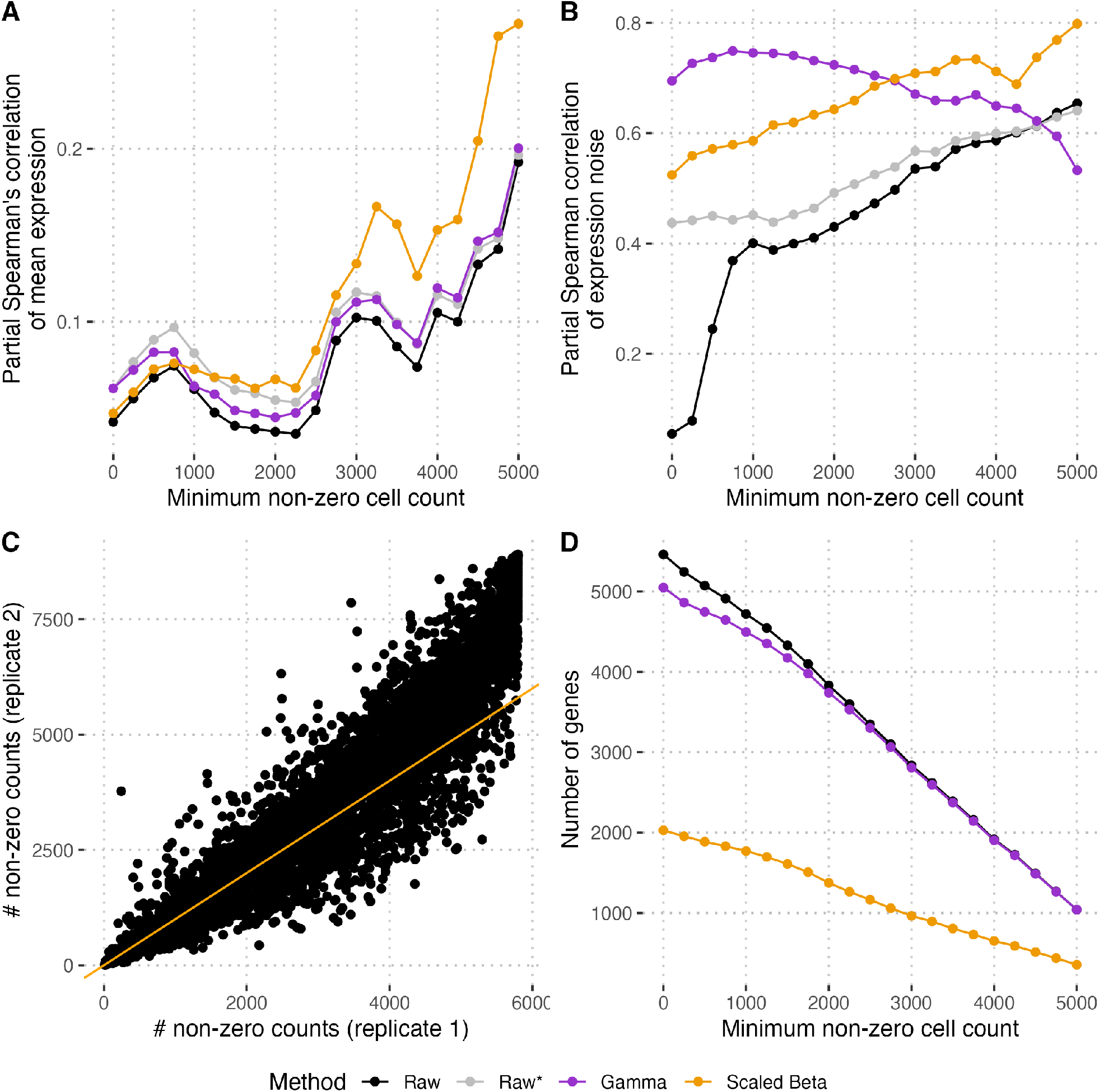
Model-based inference disentangles biological and technical noise. A) Correlations of estimates from the mean. B) Correlations of noise estimates. C) Correlation of dropout proportions. D) Number of retained genes after filtering for the maximum dropout proportion and for genes where distribution parameters could not be estimated. Partial correlations are computed using a GAM model (see Materials and methods).

### 3.4 Cell-partition models can disentangle distinct noise levels

Cell populations are characterized by different levels of heterogeneity. For example, cells are differentiated into different tissues and cell types in multicellular organisms. Unicellular organisms also show a distinct phenotype at different stages of the life cycle. This heterogeneity can complicate the inference of intrinsic expression noise. Multidimensional analysis of expression profiles, such as that performed by tools like Seurat [Hao et al., 2023], can help identify and classify cell types. We introduced a model extension that allows different cell groups to have different gene expression distributions, while sharing parameters (see Materials and methods). We apply such models to the *S. paradoxus* scRNASeq data set, using clusters identified by Seurat. In these models, cell clusters have distinct means but all share the same amount of noise. For comparison, we also fitted the same models while fixing the dropout probability to zero, as it was suggested that zero-inflation is not needed when UMI counts are modeled [sve, cao]. Whether zero-inflated or not, we found that the model with all eight clusters fitted the data best (Table 2). Interestingly, the model without zero inflation has a lower AIC when fewer clusters are modeled. However, for four clusters or more, the model with an extra dropout probability provided a better fit, despite the fact that our data are UMI counts. This could be due to a low sequencing depth or different RNA capture rates in different cell types. However, we note that dropout probabilities are inferred to be tiny, if not zero, for the majority of genes.

**Table 2:**
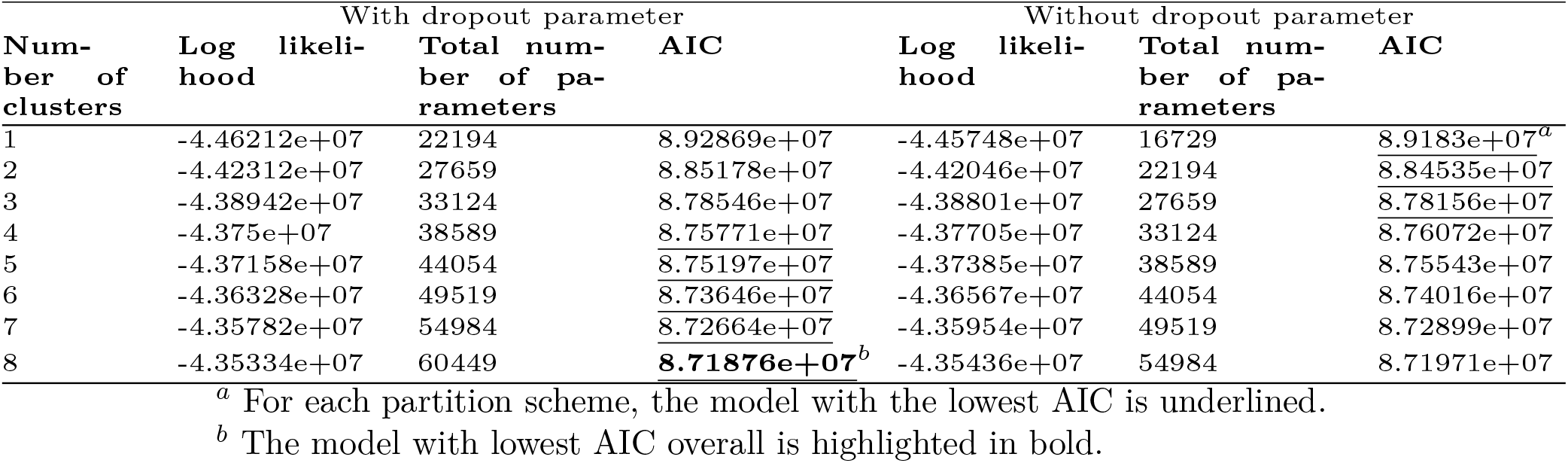
Comparison of cell-partition models fitted to *S. paradoxus* single cell data.

We conducted gene set enrichment analyzes across the models (allowing for non-zero dropout probabilities, but without filtering the resulting estimates). The analyzes revealed a consistent enrichment of protein synthesis terms and pathways for low noise genes, consistent with previous studies (Figure 4, e.g. Newman et al. [2006]). The results also revealed some terms associated with amino-acid biosynthesis that are enriched for high-noise genes. Such pathways have previously been reported to be variable and correlated in *S. cerevisiae* [Stewart-Ornstein et al., 2012]. We generally observe that accounting for cellular heterogeneity yields more significant terms and that selecting models on a per-gene basis is a conservative approach, as it retains those with the strongest significance.

**Figure 4:**
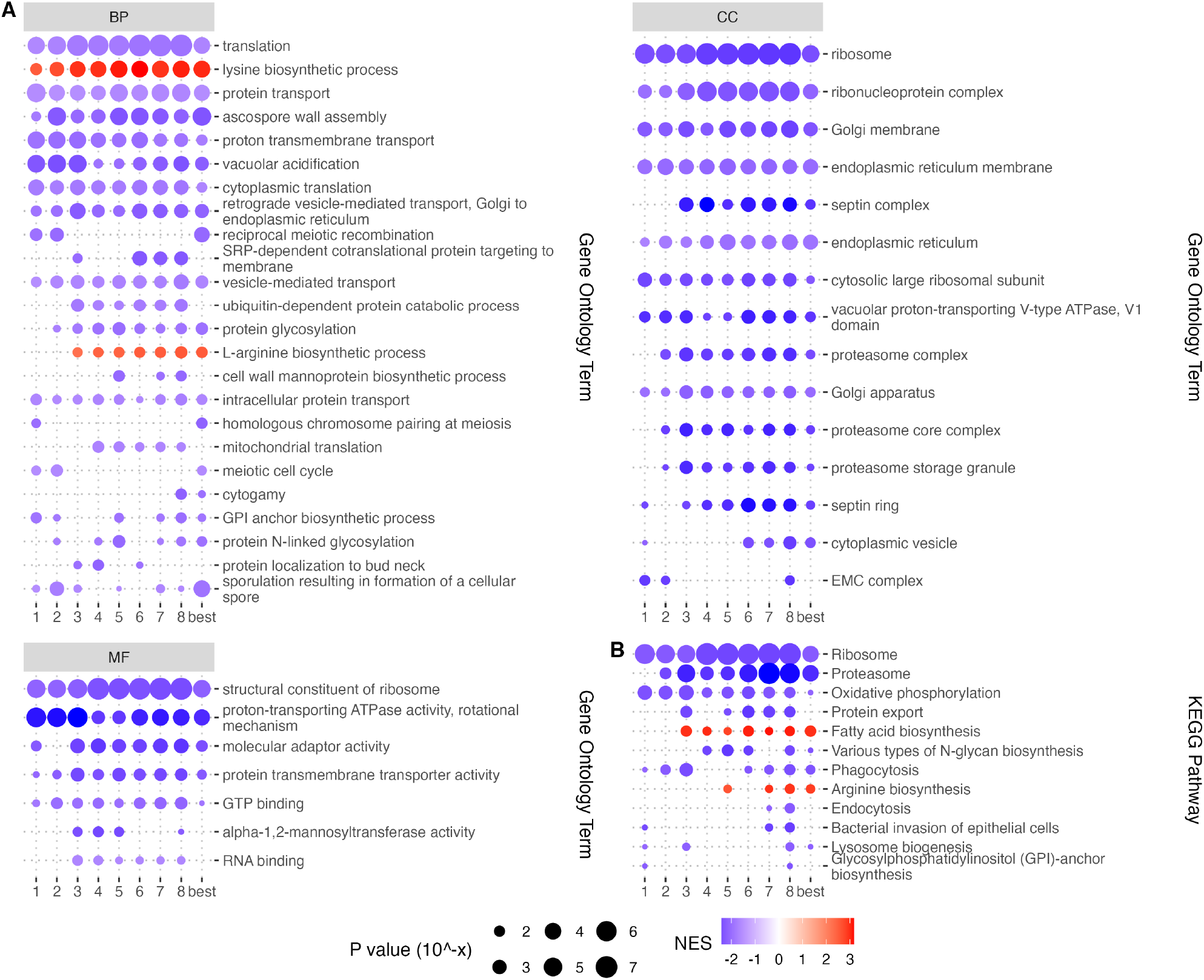
Inference of expression noise under increasingly complex cell-partition models. Genes were ranked occurring to their estimated noise value under eight distinct models: no cell clustering (*p*1), or two to eight cell clusters (*p*2*− p*8), as determined by Seurat (see Materials and methods). The column labeled as ‘best’ corresponds to the approach where the model with the lowest AIC value was selected on a per-gene basis. Significantly enriched terms at the 1% level after correction for multiple testing in at least one of the models are shown, using Gene Ontology annotation (A) and KEGG pathways (B). NES: Normalized Enrichment Score. Negative values indicate enrichment for low noise and positive values, enrichment for high noise. BP: biological process. CC: cellular component. MF: molecular function.

## 4 Conclusions

We introduced a framework for the estimation of expression noise from scRNASeq data. The framework builds on and extends standard modulated Poisson models and proposes to use the moments of the modulating distribution to estimate biological noise separately from technical noise, which is accounted for by the Poisson process. We show that the model-based estimates of noise are superior to simple estimators based on counts only. In particular, we show that the frequently observed negative correlation of expression mean and noise might be an artifact of too simple estimators. We show that Beta-Poisson models offer very good performance, but require more cells and sequencing depth than Gamma-Poisson models, which offer a good approximation for smaller data sets. When applied to real data from the yeast *S. paradoxus*, our estimates reveal relatively few terms and pathways enriched for high-noise genes. These results demonstrate that realistic models of gene expression in single cells are needed to properly assess expression noise.

## 5 Data availability

Sequencing data are available at the Sequence Read Archive, BioProject PRJNA1478450, with the sample accession numbers SRR39148149 and SRR39148148. Gene expression data are available at GEO with series accession number GSE335657, and samples accession numbers GSM9817192 and GSM9817193. The N.I.S.C.E. program is available at https://github.com/jydu/nisce. The code used to perform the analyzes can be found in https://gitlab.gwdg.de/molsysevol/expression-noise-inference.

## 6 Author contributions

**FG**: Data curation, Formal analysis, Investigation, Writing – original draft, Writing – review & editing.

**DWR**: Investigation, Writing – review & editing.

**SC**: Formal analysis, Writing – review & editing.

**JYD**: Conceptualization, Supervision, Formal analysis, Investigation, Methodology, Software, Writing – original draft, Writing – review & editing.

## 7 Acknowledgments

The experimental part of this work has been conducted in collaboration with the Competence Center for Genomic Analysis (CCGA) Kiel. The authors thank in particular Janina Fuß for sharing her expertise in short-read sequencing. The authors also thank Andrea Patricia Murillo Rincón and Markéta Kaucká for discussions on the generation and analysis of scRNASeq data.

## 10 Supplementary material

**Figure S1:**
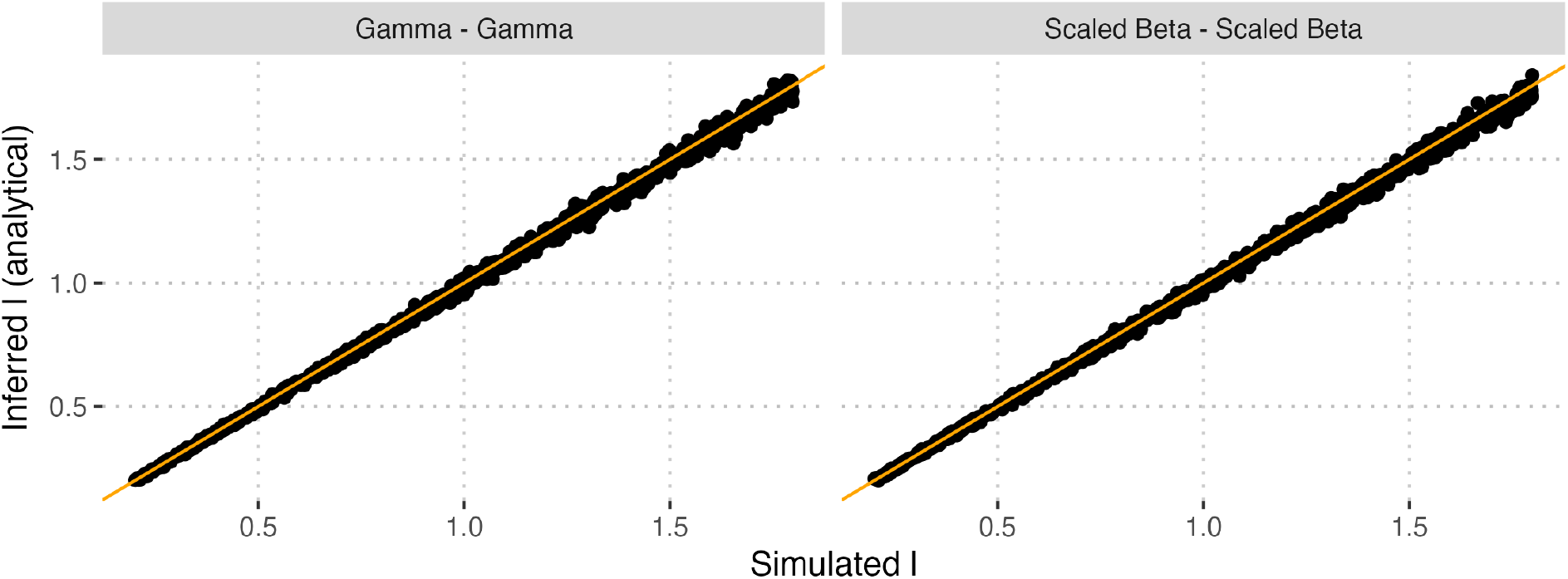
Estimation of cell normalization factors using analytical asymtotic formulas.

**Figure S2:**
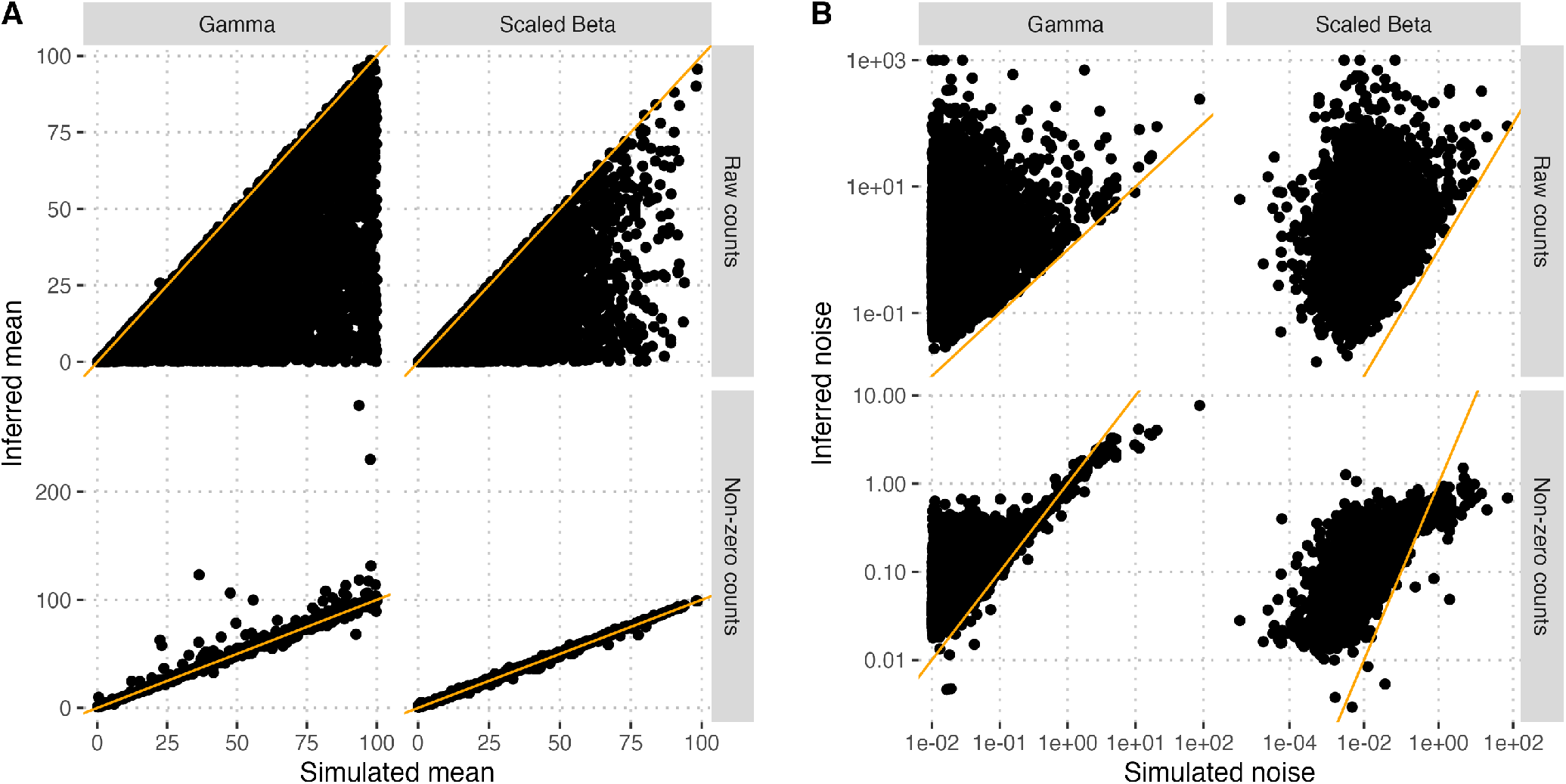
Inference of mean and noise with simple statistics. Raw-counts statistics compute the mean and noise (variance divided by the square of the mean) from the raw count data. Non-zero statistics compute the same statistics after removing all entries where counts are equal to zero.

**Figure S3:**
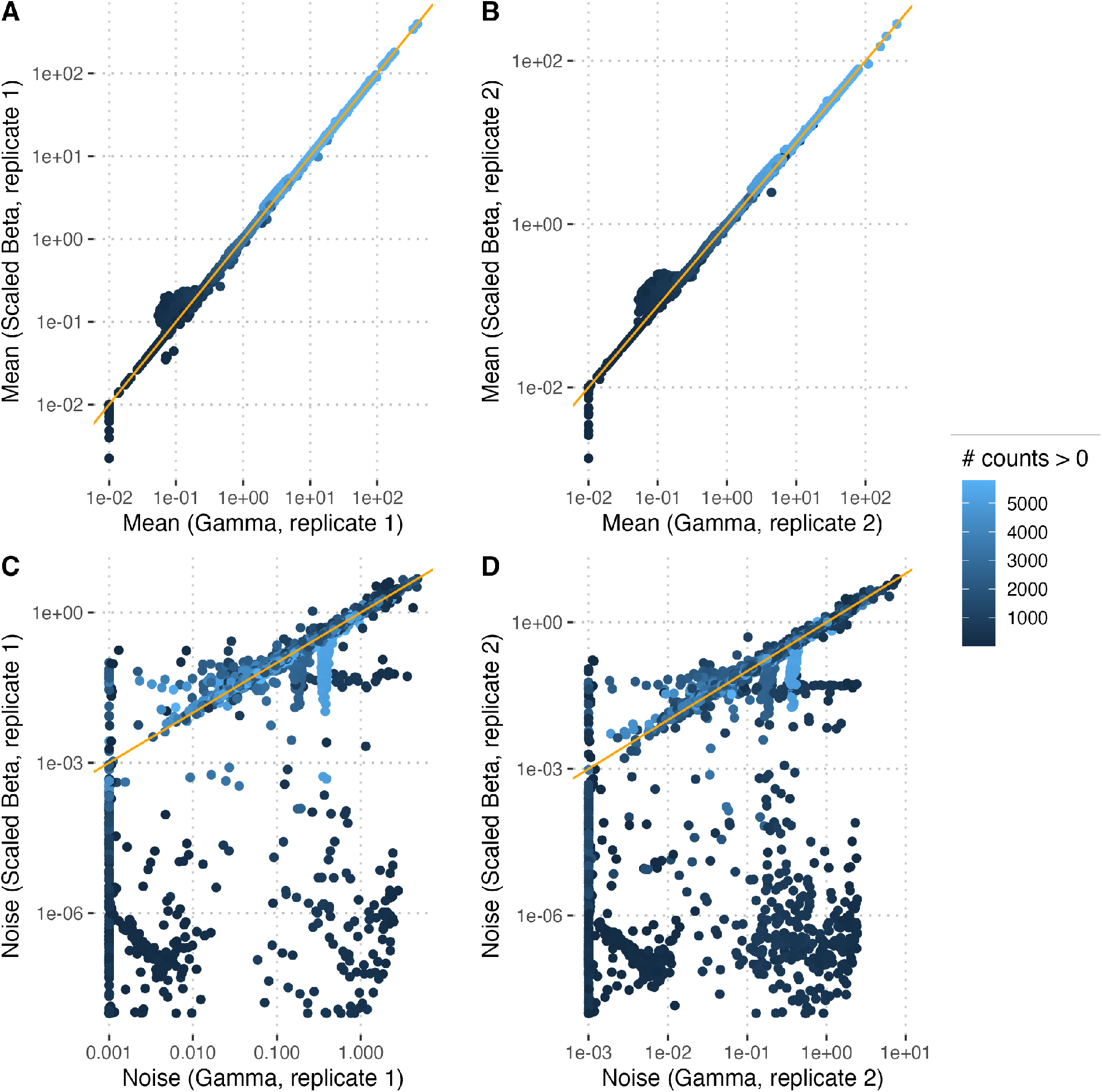
Estimation of gene expression mean and noise under a Gamma or Scale Beta model. Each point represents a gene, colored according to the number of cells with non-zero counts. The x-axis show the estimate under a Gamma model, the y-axis show the corresponding estimate under a Scaled Beta model. A and B: expression mean. C and D: expression noise. A and C: replicate 1. B and D: replicate 2.

## A Asymptotic estimates of cell normalization parameters

The expectation of number of reads in cell *j* for gene *i* is equal to

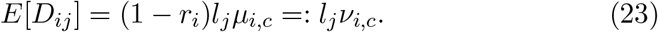

where *µ*_*i,c*_ is the mean of the distribution of mRNA for the gene in the cluster of cells *c*.

Hence, the expectation of the cell library norm is given by

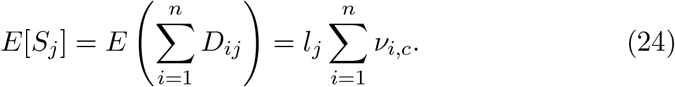

If we assume that the number of genes, *n*, is large and that a law of large numbers applies (no single gene dominates the total read count), i.e.

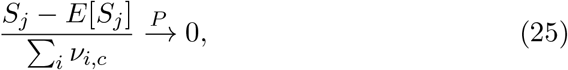

then

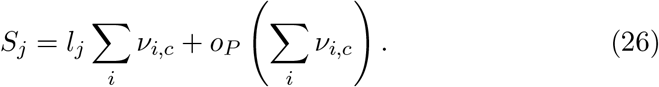

Let *C* = *{k* : *a*_*k*_ = *c}* denote the set of cells in cluster *c*, so that 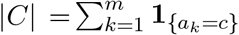. Using the identifiability constraint 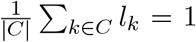, we obtain

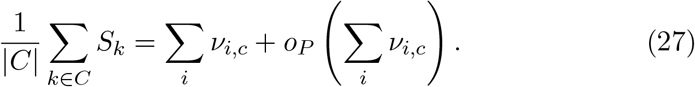

Therefore,

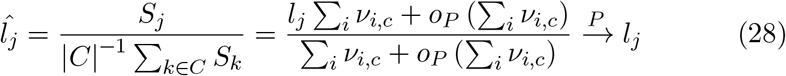

Thus, equation 17 provides a consistent estimator of the normalization constants as the number of genes increases. This provides a theoretical explanation for the close numerical agreement between the library-size estimator and the maximum likelihood estimate observed in simulations.

